# CASSPER: A Semantic Segmentation based Particle Picking Algorithm for Single Particle Cryo-Electron Microscopy

**DOI:** 10.1101/2020.01.20.912139

**Authors:** Blesson George, Anshul Assaiya, Robin Jacob Roy, Ajit Kembhavi, Radha Chauhan, Geetha Paul, Janesh Kumar, Ninan Sajeeth Philip

**Affiliations:** Artificial Intelligence Research and Intelligent Systems (airis4D), Thelliyoor - 689544, Kerala, India; Laboratory of Membrane Protein Biology, National Centre for Cell Science, NCCS Complex, S. P. Pune University Campus, Ganeshkhind, Pune-411 007, INDIA; Inter-University Centre for Astronomy and Astrophysics (IUCAA), S. P. Pune University Campus, Ganeshkhind, Pune-411 007, INDIA; Laboratory of Structural Biology, National Centre for Cell Science, NCCS Complex, S. P. Pune University Campus, Ganeshkhind, Pune-411 007, INDIA; Department of Physics, St. Thomas College, Kozhencherry - 689641, Kerala, INDIA

## Abstract

Single-particle cryo-electron microscopy has emerged as the method of choice for structure determination of proteins and protein complexes. However, particle identification and selection which is a prerequisite for achieving high-resolution still poses a major bottleneck for automating the steps of structure determination. Here, we present a generalised deep learning tool, CASSPER, for the automated detection and isolation of protein particles in transmission microscope images. This deep learning tool uses Semantic Segmentation and a collection of visually prepared training samples to capture the differences in the transmission intensities of protein, ice, carbon and other impurities found in the micrograph. CASSPER is the first method to do pixel level classification and completely eliminates the need of manual particle picking. Integration of Contrast Limited Adaptive Histogram Equalization (CLAHE) in CASSPER enables high-fidelity particle detection even in micrographs with variable ice thickness and contrast. In addition, our generalized model for cross molecule picking works with high efficiency on unseen datasets and can potentially pick particles on-the-fly, thereby, enabling automation of data processing.

## Introduction

Single-particle Cryo-electron microscopy (Cryo-EM) has revolutionized the field of structural biology by facilitating the structure determination of various biological macromolecules and their complexes^1–4^ which were recalcitrant to structure determination by X-ray crystallography or were not suitable for structure determination via NMR. Cryo-EM enables structure determination of proteins in solution without the need of protein crystallization or limitations of size, making it the current method of choice. A number of research projects are currently being carried out in hardware^5–7^ and software^8–14^ in order to streamline and automate data collection and processing steps for structure determination. One of the obstacles that still remains unresolved is the manual identification and selection of particles (protein) from micrographs for extraction and subsequent 2D classification.

To achieve high-resolution protein structure, selection of a large number of good quality particles is the prime requisite. However, particle identification, picking, and selection is a very tedious and challenging process. This is primarily due to the low Signal to Noise Ratio (SNR) of the micrographs, presence of contaminants, contrast differences owing to varying ice thickness, absence of well segregated particles etc. To overcome these drawbacks introduced by EM grid vitrification and low dose imaging, one often has to rely on laborious, slow, manual or semi-automated methods. A fast, automatic method that can replace the manual processing is thus a necessity for automating the structure determination process.

Presently, considerable effort is being devoted to the development of automated particle picking methods in order to circumvent the manual intervention. These can be broadly categorized into two groups (i) Template free and (ii) Template based methods which rely mainly on cross correlation with the template images. Gautomatch^15, 16^ is one of the widely used template free methods based on cross correlation. In RELION^11^ and cryoSPARC^12^, a Gaussian blob of defined size is used as a template for particle picking. Similarly, DoGpicker^17^ uses mathematically derived Gaussian functions as templates to recognize and select particles from the micrographs. However, these tools are prone to picking huge amount of contaminants, background, and ice, and do not work optimally for datasets with poor SNR or small particle sizes. These problems are resolved to a certain extent in template (reference) based particle picking tools implemented in SIGNATURE^18^, RELION^11^, cryoSPARC^12^, EMAN^8^, SPHIRE^13^, cisTEM^19^, FindEM^16^, gEMpicker^20^ etc. In all these methods, templates are generated by manually picking a few hundred to several thousand particles from multiple micrographs. These particles are then sorted and 2D classified to generate templates for automated particle selection via template matching algorithms. While this methodology works better than previously described reference-free methods, it is time consuming, computationally expensive, and also requires manual intervention preventing its integration into automated pipelines for structure determination. In addition, manual particle picking usually introduces a strong template bias that may result in high false picking rate.

Artificial Intelligence/Machine Learning (AI/ML) based approaches have the potential to overcome the problems discussed above and pave the way for full-automation of the data processing pipeline. Not surprisingly, multiple AI/ML based methods have also been proposed, such as, XMIPP^21^, APPLE picker^22^, DeepPicker^23^, DeepEM^24^, FastParticle Picker^25^, crYOLO^26^, PIXER^27^, PARSED^28^ etc that are based on Convolutional Neural Networks (CNN), Region-based Convolutional Neural Networks (R-CNN), cross correlation, and segmentation respectively. These deep learning classifiers are first trained on available datasets with known labels. The training process allows them to learn intrinsic and unique features/characteristics of the particles. A trained classifier can later be used to automatically pick similar particles from other micrographs. In Convolutional Neural Network (CNN) based methods like DeepPicker and DeepEM, a sliding window is used to analyze the image for classification. In crYOLO, which is also a CNN based tool, the entire image is split into grids and part of the image in each grid is taken as an input to the classifier. However, all the above mentioned methods require individual particles to be manually picked for training, which is a time consuming procedure. Further, the exposure difference, noise level and the variable ice thickness in micrographs also limits the performance of these tools.

Here, we present a novel method packaged as “CASSPER” (**C**ryo-EM **A**utomatic **S**emantic **S**egmentation based **P**article pick**ER**) based on Semantic Segmentation (SS) for automated particle picking with high precision and accuracy. To our knowledge, CASSPER is the first method to carry out labelling and prediction of different kinds of particles (protein, ice, carbon etc) in a micrograph. Employing SS, CASSPER learns how to differentiate each pixel of the image by considering the transmittance of the medium. Since protein, ice and carbon contamination differ in transmittance, CASSPER can differentiate between them and provide unique labels with high confidence and reliability. CASSPER, like other AI based techniques, requires a training data. However, it has a Graphical User Interface (GUI) with a few sliding bars that can be used to label all the particles in a micrograph in one go, making it highly efficient and time saving in the preparation of the training data. Further, CASSPER utilizes the Contrast Limited Adaptive Histogram Equalization (CLAHE)^29^ algorithm for efficient particle identification even in micrographs with large regional contrast variability. Since CASSPER learns the pixel differences and not the particle morphologies, it yields high accuracy even on unseen micrographs.

## Results

### Implementation of Semantic Segmentation in CASSPER

Unlike the traditional image classification methods that use either derived features or morphological characteristics of the target image for its identification, CASSPER uses Semantic Segmentation (SS) for identifying the protein molecule at the pixel level^30^. CASSPER is coded in the Python language and the Deep Learning model uses InceptionV4^31^ for feature extraction and a Full Resolution Residual Network (FRRN)^32^ model for SS. The FRRN has two different processing streams; namely, a full resolution residual stream that holds high resolution details for recovering the location of the detections and a pooling stream for extracting the hidden features required for learning the abstract relationships in the image. By using a set of Full Resolution Residual Units (FRRU) to merge the residual stream and the information from the pooling layers at each stage, it ensures localisation as well as classification accuracy during reconstruction. This is one advantage of using FRRN instead of the more popular Convolution Neural Network (CNN). Each FRRU is a combination of Residual Units (RU) and Feed Forward Networks, where the latter are built using a linear sequence of layers. The output of the *n^th^* layer of the Feed Forward Network is given by *y*_*n*_ = *f*(*y_n_*_-_*_1_*; *W*_n_) and can be represented as a function of output of (*n* − *1*)*^th^* layer and the *W_n_* parameters of the layer. The Residual Network^33^ is composed of a series of RUs and for them, the output of the *n^th^* layer is given as *y_n_* = *y_n_*_-_*_1_* + *f*(*y_n_*_-_*_1_*; *W_n_*), where*f*(*y_n_*_-_*_1_*; *W_n_*) is the Residual of the layer. The outputs of the *n^th^* layer FRRU is given by:

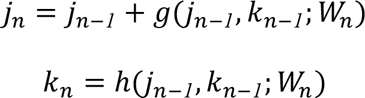

Where *j_n_*_-_*_1_* and *k_n_*_-_*_1_* are the residual and pooling inputs to FRRU. The FRRN architecture that is used for the present study employs 5 FRRUs for upsampling and 4 FRRUs for downsampling.

Also it has 5 maxpooling and 4 unpooling layers in the pooling stream. The network design is explained in **Figure 1** **(A).** CASSPER uses the SS implementation by George Seif^34^.

**Figure 1.**
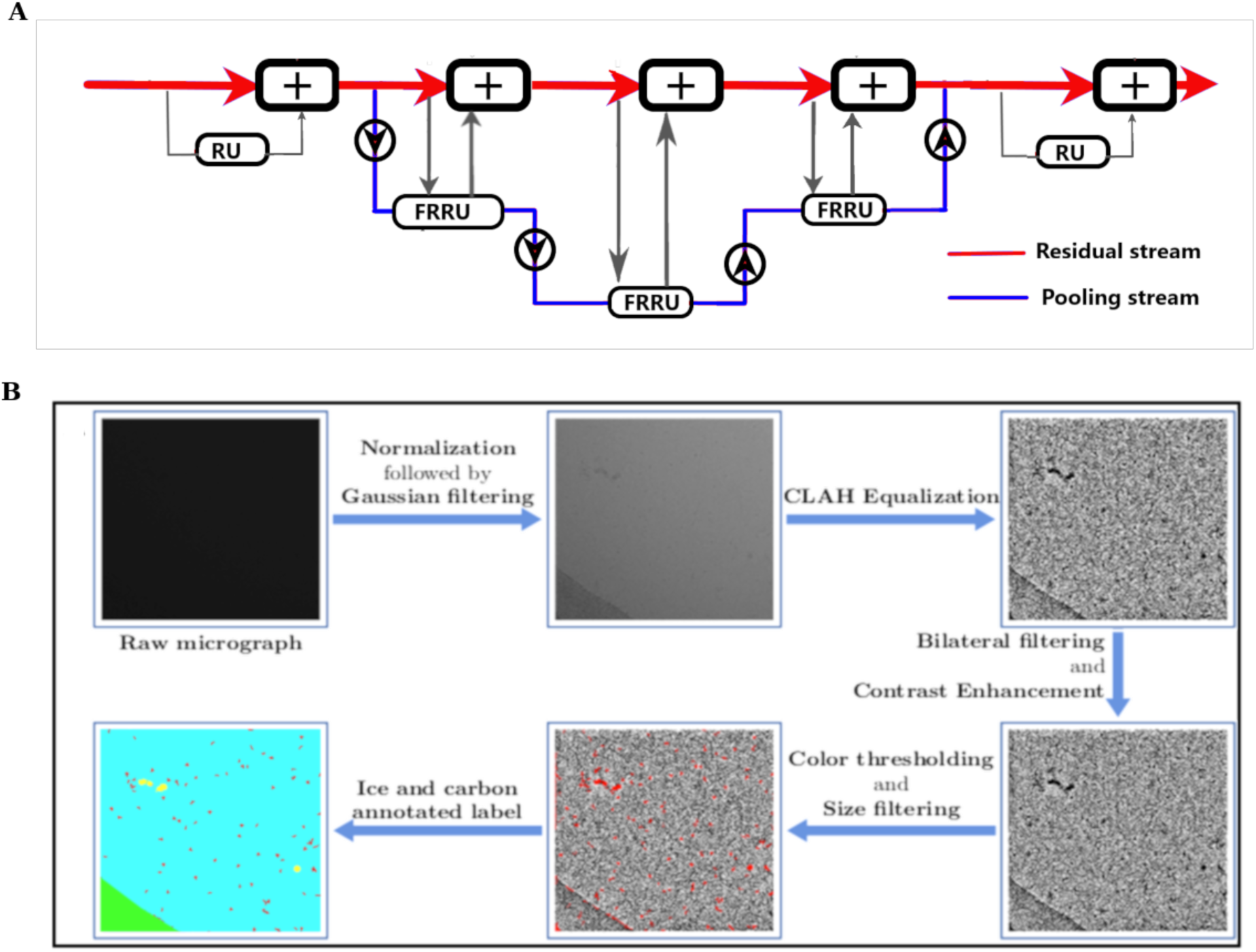
**(A) Abstract structure of Full Resolution Residual Network (FRRN).** FRRN achieves better recognition and localization performance by image processing in two different streams; namely, pooling and residual streams. Pooling stream learns the abstract relationships in the image and residual stream carries a full resolution feature map that ensure localization capability. **(B) Flowchart of the Image labelling tool which is used for preparing training data for Semantic Segmentation.** The motion corrected mrc is filtered using Gaussian filter after normalization. This step makes the micrograph more visible for the user. Contrast Limited Adaptive Histogram Equalization (CLAHE) is applied to eliminate any exposure differences in the image and bilateral filtering followed by contrast enhancement makes the difference between background and particles more vivid. Intensity thresholding is done to distinguish particles from background. Size thresholding is done for removal of contaminants or background that is not eliminated at intensity thresholding. The selected parts of the mrc are indicated in red. The ice and carbon contaminations are later labelled selecting the regions of interest.

### CASSPER pipeline and preparation of training data

The entire pipeline of CASSPER can be divided into two phases - Training phase and Prediction phase. Like all other supervised Machine Learning methods^35^, SS also requires a training data. The training data is prepared by labeling the different objects of interest in the image by assigning them different colours. Thus, the label itself is an image with coloured pixel masks indicating their type. Since it is a pixel based learning method, the accurate labeling of each pixel is crucial. In order to carry out labelling with minimum user intervention, we developed a graphical labelling tool. The tool enables visual enhancements in the image by varying its contrast, bilateral filter size, intensity and threshold values. All controls are implemented with the help of slide bars explained in the methods. A schematic of the functionalities of the slide bars are shown in **Figure 1** **(B)**. The method is independent of the structural details of the protein and hence its efficiency is unaltered by the differences in shape or size of the projected image of the protein. An illustration of the same is shown in **Figure 2** where the micrographs are labeled to show four different constituents (referred to as classes hereafter), namely crystalline ice, carbon edges, background and the protein molecules. About 15-25 micrographs of each protein were labelled and used to train the network. The raw and labelled micrographs for training were provided with the same root names in the pipeline and about 90% of them were cycled in ∼300 epochs to train the network. The remaining 10% of the data was used for validating the performance of the trained network. The training round that gives best F1 scores (see section Statistics for details) during validation was taken as the criterion for choosing the final trained model. Subsequent prediction on the larger set of unlabelled micrographs was done using the trained model. F1 scores for a few proteins during validation are shown in **Figure 3** **(A).** CASSPER labels each particle with the same colours that were used to represent them in the training data. Since we are interested only in finding the coordinates of the protein, everything other than the protein is masked out from the image. The user is then allowed to specify the size of a circular mask approximately the size of protein. The reason to take it as an input is that, occasionally, the image of the protein may appear fragmented and the machine needs this information to include those fragments as part of the same particle. CASSPER then estimates the centers of those contours and its coordinates are provided in the “star format” for particle extraction and subsequent processing steps.

**Figure 2.**
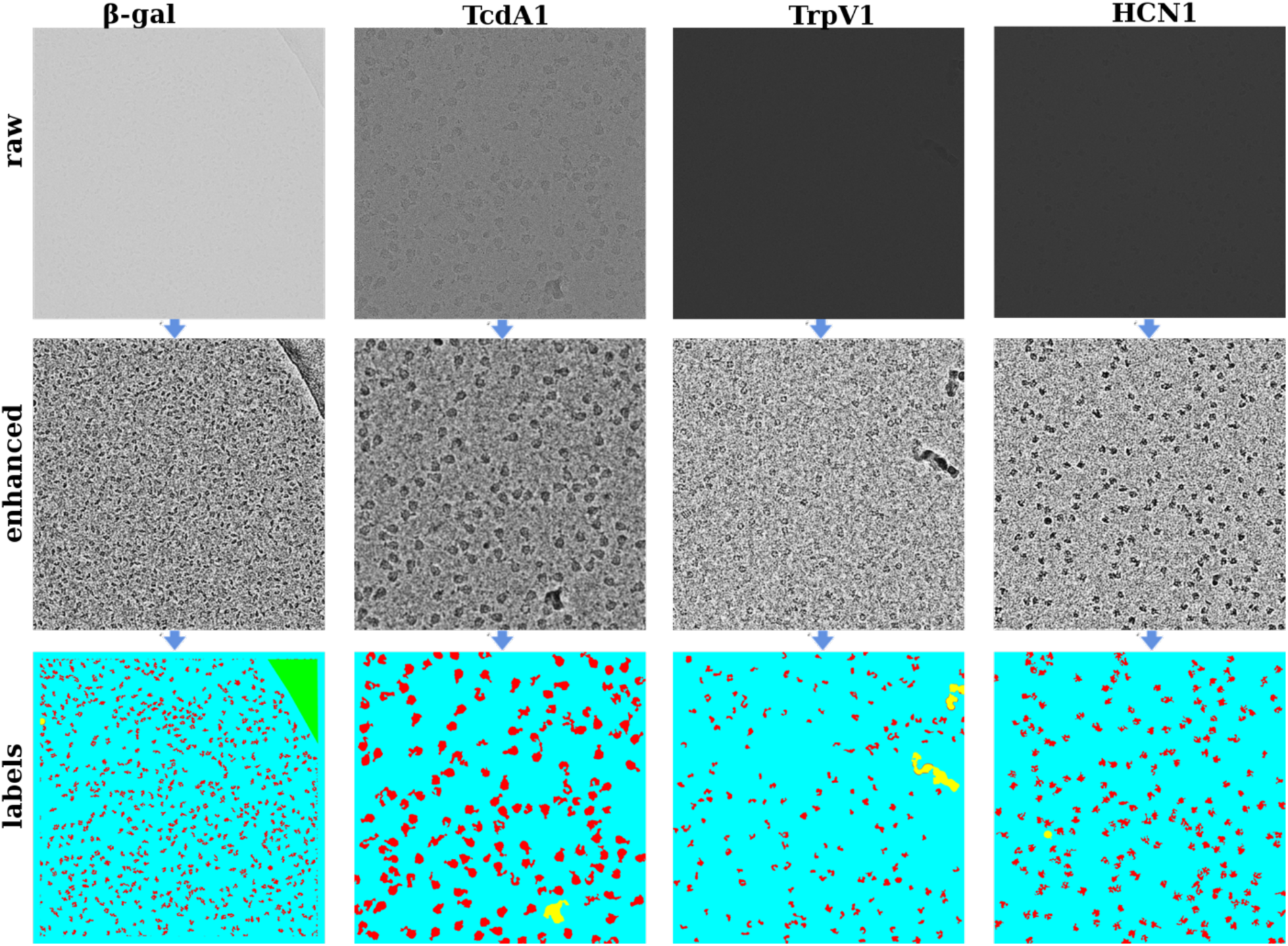
The raw micrographs, contrast enhanced and Semantically Segmented images of micrographs from β-galactosidase (EMPIAR 10017), TcdA1 (EMPIAR 10189), TRPV1 (EMPIAR 10005) and HCN1 (EMPIAR 10081) respectively. The entire micrograph is segmented into different classes and each class is represented by a unique color. The background, protein, ice contamination and carbon contamination are represented using cyan, red, yellow and green color respectively.

**Figure 3.**
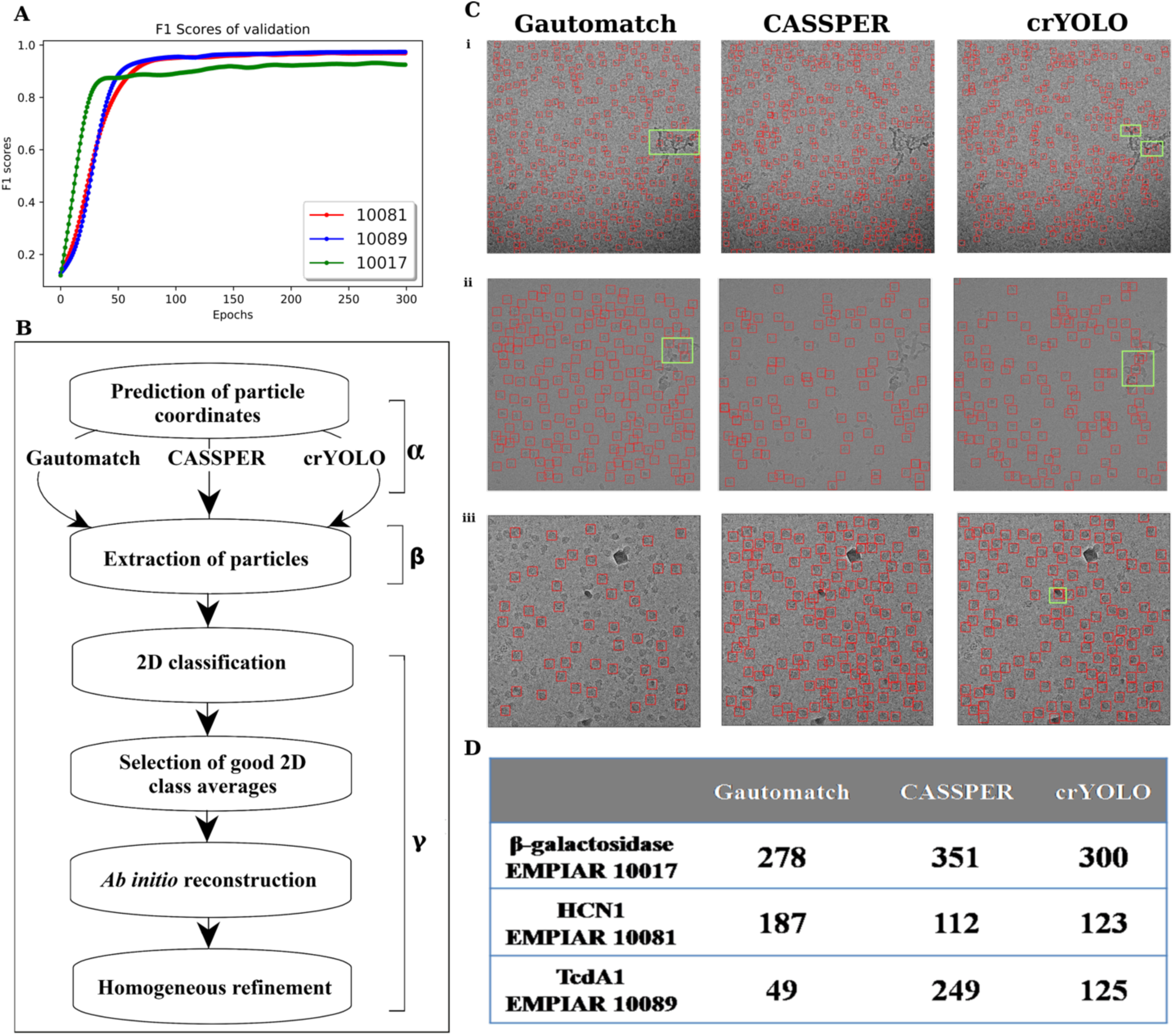
**(A)** The validation F1 scores, which are used to select the best trained model in training epochs for prediction, for all three proteins. **(B)** Schematic representation of the uniform pipeline used to compare the performance of Gautomatch, CASSPER and crYOLO. **(C)** Representative micrographs for (i) β-galactosidase, (ii) HCN1 and (iii) TcdA1 showing the particle picking performance of different tools. Highlighted areas indicate the noise picked by the respective tools. **(D)** Table showing the number of particles picked by Gautomatch, CASSPER and crYOLO on same micrograph for each protein as shown in panel (C).

The evaluation of the performance of CASSPER was carried out in three steps:

1. The number of good 2D classes and the number of particles therein.
2. The resolution of 3D map generated using a uniform processing pipeline as described below.
3. F1 and IoU scores for labels created by CASSPER on randomly selected micrograph were compared with ground truth values.

### Uniform pipeline for comparison

The performance of CASSPER was evaluated by comparing the results with two popular particle picking tools - machine learning based crYOLO and cross correlation based Gautomatch. A uniform pipeline for cryo EM data processing was adopted to rationally assess the quality of results from the three methods over a limited number of steps. The uniform pipeline scheme can be divided into three phases illustrated in **Figure 3** **(B).** Phase **α** predicts the particle coordinates using Gautomatch, CASSPER and crYOLO. The same micrographs were used for particle prediction by all the three tools. Comparison for the true positive (actual protein particles) and false positive (wrong prediction as protein) particles on one representative micrograph for each protein is depicted by **Figure 3(C)** and the comparison has been summarized in **Figure 3(D)**. In this phase, each tool outputs a star file with the particle coordinates they identify. In Phase **β**, the particle coordinates were imported into RELION^11^ and the particles were extracted from the CTF estimated micrographs. The extraction box size is kept uniform for each protein predicted with different tools. Phase **γ** of the pipeline is carried out in cryoSPARC v1 to accelerate the data processing steps. The extracted particle stack obtained by the three tools for a sample of four datasets (explained in the next section) were imported into cryoSPARC v1 where 2D classification was performed. After a single round of 2D classification, class averages with discernible features were selected and used for *ab initio* reconstruction with C1 symmetry. Later the 3D models generated from them were refined with single step homogenous refinement in their respective symmetry groups. The number of good 2D classes and particles therein and the resolution of the map in subsequent 3D reconstruction achieved indirectly reflects the performance of the particle picking tools (**Table 1**).

**Table 1.** Comparison of Gautomatch, CASSPER and crYOLO for β-galactosidase, TcdA1, TRPV1 and HCN1. The total number of particles picked by the respective tools were fed into the uniform pipeline scheme for further processing. The 2D class averages with characteristic features were selected and used for 3D reconstruction followed by homogeneous refinement for all proteins by imposing the respective symmetry. The resolution obtained through uniform pipeline scheme are given in the table.

| **Protein** | **Method** | **No of particles picked** | **No of particles selected for 3D reconstruction** | **Resolution ($\text{\AA}$)** |
| --- | --- | --- | --- | --- |
| **TcdA1**<br>EMPIAR 10089 | GAUTOMATCH | 4097 | 3364 | 6.7 |
|  | CASSPER | 14603 | 11245 | **4.3** |
|  | crYOLO | 11127 | 10629 | 4.4 |
| **β-gal**<br>EMPIAR 10017 | GAUTOMATCH | 25409 | 21476 | 10.8 |
|  | CASSPER | 44261 | 40467 | **7.26** |
|  | crYOLO | 44591 | 42876 | 7.32 |
| **HCN1**<br>EMPIAR 10081 | GAUTOMATCH | 195782 | 107332 | 5.31 |
|  | CASSPER | 150342 | 115297 | **5.06** |
|  | crYOLO | 141002 | 103010 | 6.02 |
| **TRPV1**<br>EMPIAR 10005 | GAUTOMATCH | 127776 | 38836 | 7.73 |
|  | CASSPER | 107320 | 46913 | 3.74 |
|  | crYOLO | 110153 | 67269 | **3.68** |

We trained crYOLO for each dataset and the trained model was used for comparison. In the case of Gautomatch, pixel size and particle diameter were the only two parameters which were used to predict the particles. (./Gautomatch-{version} --apixM{pixel size} --diameter {particle diameter} /path of the folder containing micrographs).

### CASSPER Performance

To test the performance of CASSPER, we selected four datasets, namely; HCN1^36^ (EMPIAR 10081), TRPV1^37^ (EMPIAR 10005) TcdA1^13, 26^ (EMPIAR 10089) and β-galactosidase^38^(EMPIAR 10017). Our selection includes proteins with different molecular weight (464kDa - 1.4 MDa) and proteins from different biological environments ranging from cytoplasmic to membrane proteins.

### TcdA1

TcdA1 (EMPIAR 10089) is one of the well studied components of tripartite ABC type toxin complexes released by nematodes in case of insect invasion. It has a molecular weight of 1.4 MDa and its characteristic shape renders it clearly distinguishable on the micrographs making it a suitable candidate to develop an autopicking tool. This dataset was obtained from the EMPIAR database and has 97 movies which were acquired on Titan Krios with Falcon II detector (4k x 4k). The raw movies were motion corrected by MotionCorr2^39^, and CTF estimation was performed using CTFFIND4^40^ in RELION. 26 micrographs were randomly picked and labelled for training *via* CASSPER. The actual training used 23 micrographs and 3 micrographs were used for validation. The different validation parameters were monitored in all epochs. The maximum F1 score and mean IoU score obtained for the training data were 0.98 and 0.88 respectively. The SS model with highest F1 score as indicated in **Figure 3** **(A)** was used for making the predictions. The coordinates of the centers of the predicted particles were returned in star format. CASSPER showed the best performance for TcdA1 as it produced 11245 good particles for the 3D reconstruction from 97 micrographs **(Table1)**.

### β-galactosidase

β-galactosidase (EMPIAR 10017) is a soluble protein which is routinely used for benchmarking cryoEM data processing tools and softwares. It is derived from *E.coli* and forms a biological tetramer whose molecular weight corresponds to 464 kDa. The data was obtained from the EMPIAR database and was acquired by POLARA with a Falcon II detector. For the study, 84 micrographs were used. The maximum F1 score and mean IoU score for training were 0.93 and 0.75 respectively. Out of 44261 particles picked from 84 micrographs by CASSPER, 40467 were used for 3D reconstruction. Homogeneous refinement was performed by enforcing D2 symmetry for the 3D maps to obtain a resolution of 7.26 Å which is better than the other tools under uniform pipeline approach **(Table1)**.

### HCN1

HCN1 (EMPIAR 10081) is also a membrane protein which plays a pivotal role in controlling the rhythmic activity of cardiac and neuronal cells. A total of 997 micrographs obtained from the EMPIAR database were used by all the three tools to predict the protein particles. After the 2D classification step in the uniform pipeline, 76% of the total number of particles picked by CASSPER were used for 3D reconstruction. It must be noted that CASSPER picked nearly 8000 more true positive particles than the other tools used in this study. The difference in number of particles picked by these three tools for 3D reconstruction through the uniform pipeline corresponding to the difference in their resolution are shown in **Table1**.

### TRPV1

TRPV1 (EMPIAR 10005) is involved in mediating response to various physical and chemical stimuli from the environment. For this dataset, 771 micrographs obtained from the EMPIAR database (collected on FEI POLARA 300 using GATAN K2 detector) were used to compare the performance of CASSPER with other tools through the uniform pipeline approach. Even though the total number of particles picked by CASSPER is less, 2D class averages and 3D maps are comparable with other tools.

The 2D class averages for all the datasets obtained by processing the coordinates from different tools showed similar features (**Figure 4**). However, the final 3D maps after refinement clearly indicate the difference in the resolution. The EM density maps for β-galactosidase, TcdA1, TRPV1 and HCN1 are shown in **Figure 5** **(A)**. Resolutions for all the EM density maps were estimated by Fourier shell correlation at FSC= 0.143 criterion indicated in **Figure 5** **(B-E)**

**Figure 4.**
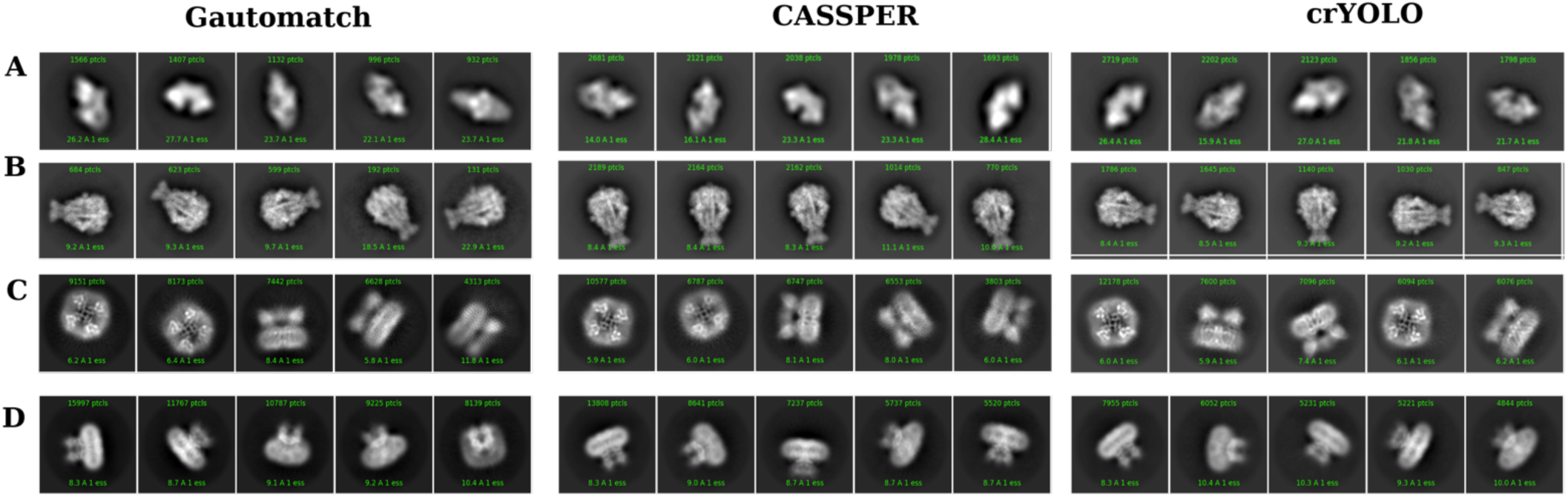
**Comparison of Representative 2D class averages** for **(A)** β-galactosidase, **(B)** TcdA1, **(C)** TRPV1 and **(D)** HCN1 obtained after single round of 2D classification in uniform pipeline using the particles picked by different tools.

**Figure 5.**
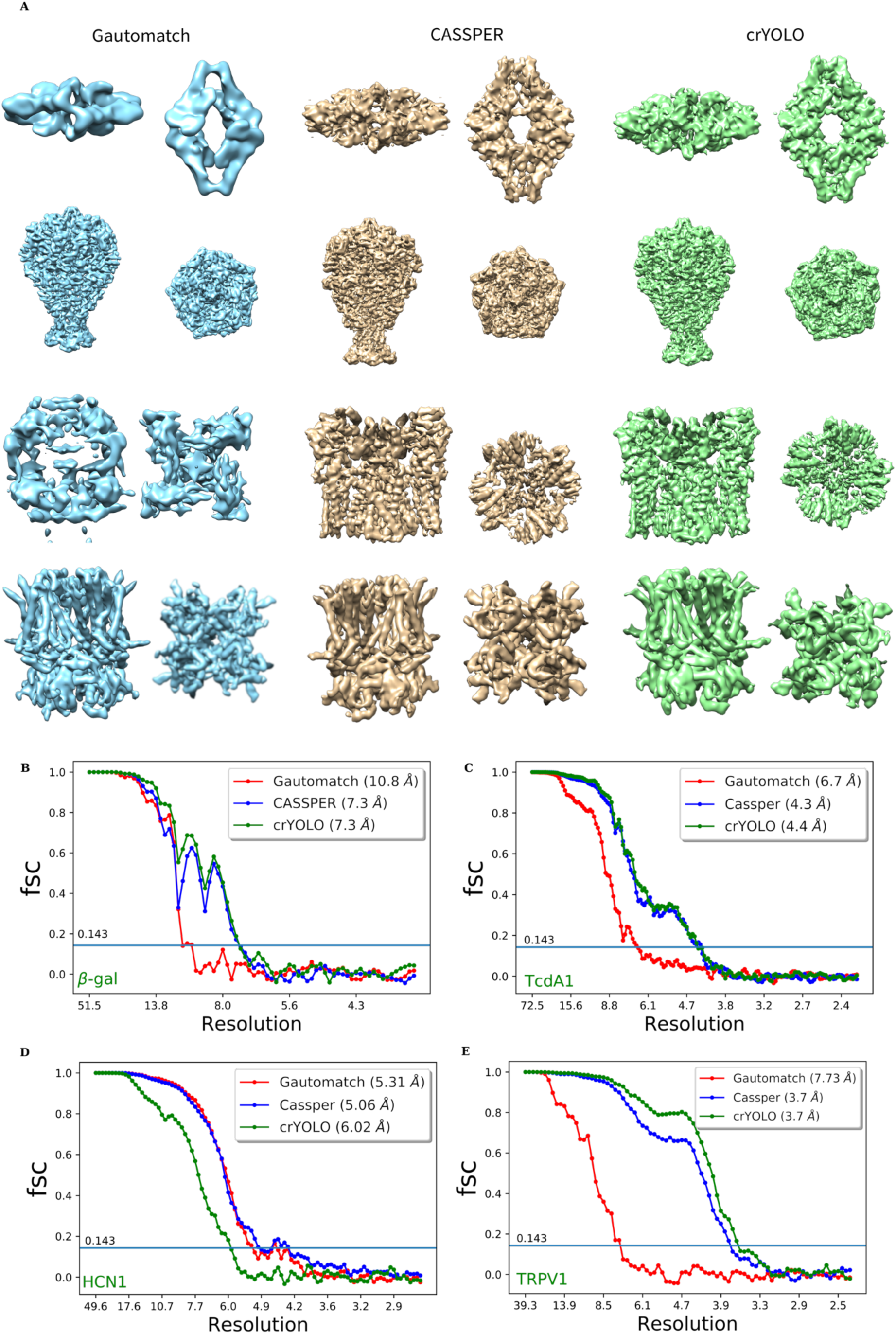
Comparison of 3D models for β-galactosidase, TcdA1, TRPV1 and HCN1 generated using particles picked by Gautomatch (blue), CASSPER (tan), crYOLO (green). The particles were extracted in RELION2 and further processing was done using cryoSPARC v1 as per the uniform pipeline scheme. **(A)** Different views of the 3D models generated for β-galactosidase, TcdA1, TRPV1, and HCN1. FSC curves (tight mask) for the 3D reconstruction of β-galactosidase **(B)**, TcdA1 **(C)**, TRPV1 **(D)** and HCN1 **(E)** showing the resolution at the gold standard cut off (0.143) obtained using Gautomatch (red), CASSPER(blue) and crYOLO(Green)

### Evaluation of the predicted labels

The labels generated in the training step using the CASSPER labelling tool were treated as the ground truth labels to evaluate the quality of the predicted labels. 12 micrographs which do not constitute the training dataset were randomly selected for each protein and mean values of F1, IoU and accuracy scores, shown in **Figure 6**, were obtained by comparing the predicted labels with the corresponding ground truth. The high scores of evaluation metrics comprised of mean F1, IoU and accuracy scores suggest that our model is robust and performs proficiently irrespective of varying protein sizes and contrast differences across the micrographs.

**Figure 6.**
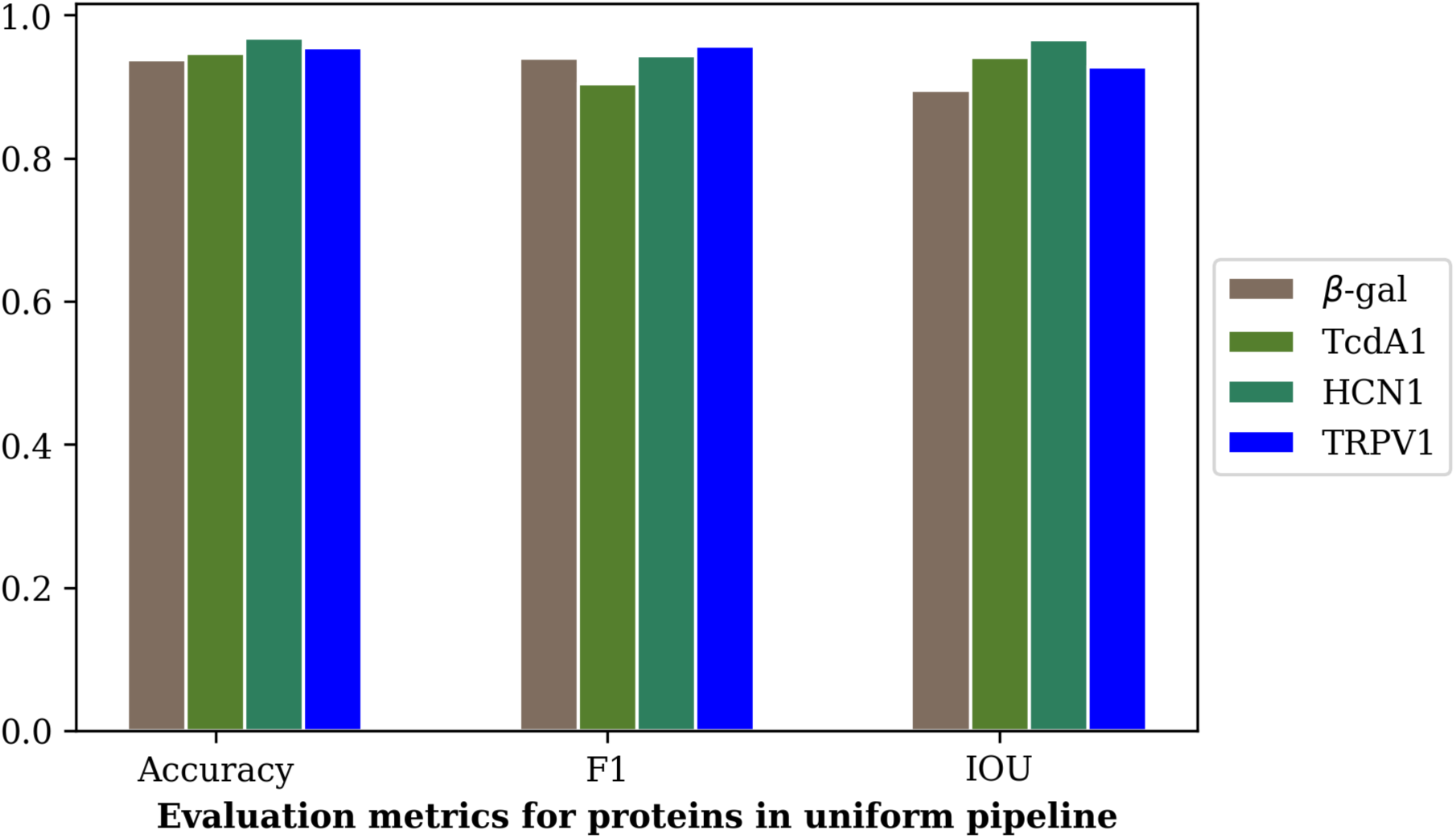
The mean values of accuracy, F1 and mean IoU scores of proteins employed in uniform pipeline obtained by comparing the labels predicted using trained SS and ground truth labels are shown. The high values of these scores show that the protein particles are accurately predicted by the trained model.

### High-resolution 3D reconstruction

The ultimate goal of macromolecular structure determination is to explore biologically relevant intramolecular and intermolecular interactions in its native environment. This is possible only if we achieve high-resolution structure which furnishes atomic level details. To demonstrate the utility of CASSPER in obtaining high-resolution 3D reconstruction, we processed cryo-EM data for TcdA1 and TRPV1 datasets following the homogeneous, non-uniform and local refinement protocols implemented in cryoSPARC v2 and obtained a resolution of 3.5 Å and 3.19 Å respectively. The resolution for TcdA1 obtained with CASSPER is equal to the previous report, however for TrpV1 the resolution with CASSPER was better than that of the published report.^13, 37^

**Figure 7** **and** 8 show the final 3D maps of TcdA1 and TRPV1 obtained using the coordinates derived from CASSPER where high-resolution features are clearly visible. This clearly demonstrates the ability of CASSPER to automatically pick high-quality particles for high-resolution structure determination.

**Figure 7.**
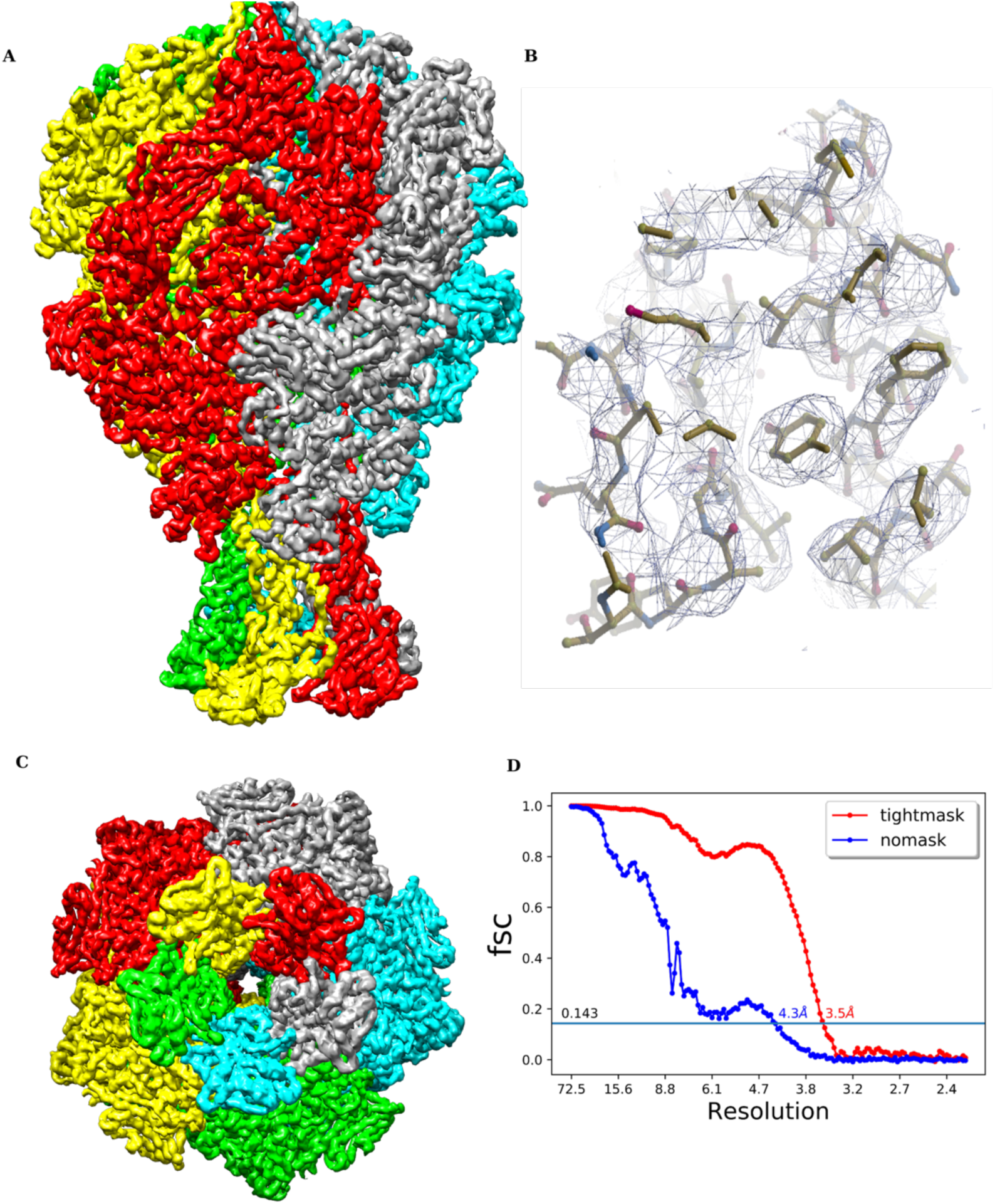
3D EM density map of TcdA1 obtained by implementing additional refinement steps to the uniform pipeline scheme. **(A)** Side view **(B)** Highlighted view of the side chains fitted (PDB 1VW1) into the EM density **(C)** Top view **(D)** FSC curve for TcdA1 showing resolution (Å) at gold standard cutoff (0.143).

**Figure 8.**
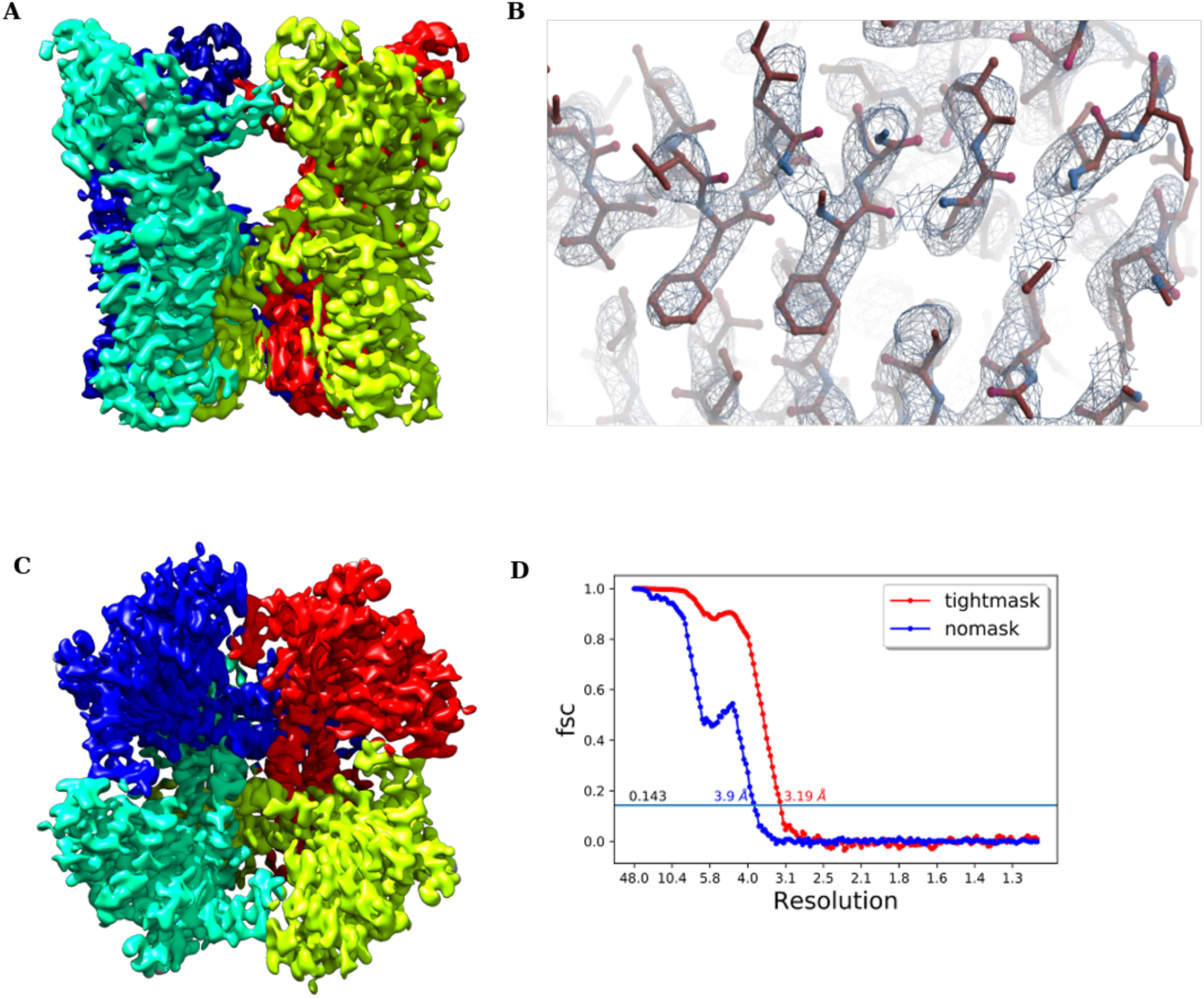
3D cryo EM density of TRPV1 obtained by implementing additional refinement steps to uniform pipeline scheme. **(a)** Side view **(b)** Highlighted view of the side chains fitted (PDB 3j5p) into the EM density **(c)** Top view **(d)** FSC curve for TRPV1 showing resolution (Å) at gold standard cutoff (0.143).

### Analysis of Generalization ability

The ability of SS method to learn the intrinsic features and composition of different objects along with their context was used to implement a cross protein classification model to predict the particle coordinates in unseen micrographs. The cross model was trained on 74 micrographs from six datasets (HCN1, TRPV1, β-galactosidase, RNA Polymerase III (EMPIAR 10168), Influenza Hemagglutinin (EMPIAR 10096), T20S proteasome (EMPIAR 10057)) and predicted for TcdA1. The cross model predicted 13209 particles from 97 micrographs of TcdA1 which after undergoing data processing steps in uniform pipeline resulted in 3D reconstruction with 4.38 Å resolution which is comparable with the CASSPER trained model under uniform pipeline scheme. Similarly the prediction for HCN1 based on cross model training gave a resolution of 5.7 Å.

In an attempt to improve the generalization ability of the cross model, we developed a cross CASSPER model trained with 180 micrographs from fifteen different proteins. To analyze the generalization ability of the cross model, 12 micrographs each from **HIV-1 envelope glycoprotein** (EMPIAR 10004), **T20S proteasome** (EMPIAR 10057), **80S ribosome** (EMPIAR 10028) and **afTMEM16/nanodisc complex** (EMPIAR 10240) were predicted using the cross model. These four proteins were totally new and unseen by CASSPER during the training phase. The predicted images were compared with the ground truth labels prepared using CASSPER labelling tool. Different metrics such as F1, Accuracy and mean IoU scores were used to evaluate the performance of the predicted labels. The representative images of the predicted micrographs along with box plot showing Accuracy, Precision, F1 and Mean IoU scores are given in **Figure 9**. The plots, with more than 90% scores, ascertain that our cross CASSPER model performs very efficiently in predicting the particles even for the unseen datasets, making it suitable for integration in any of the available cryo-EM data processing pipelines.

**Figure 9.**
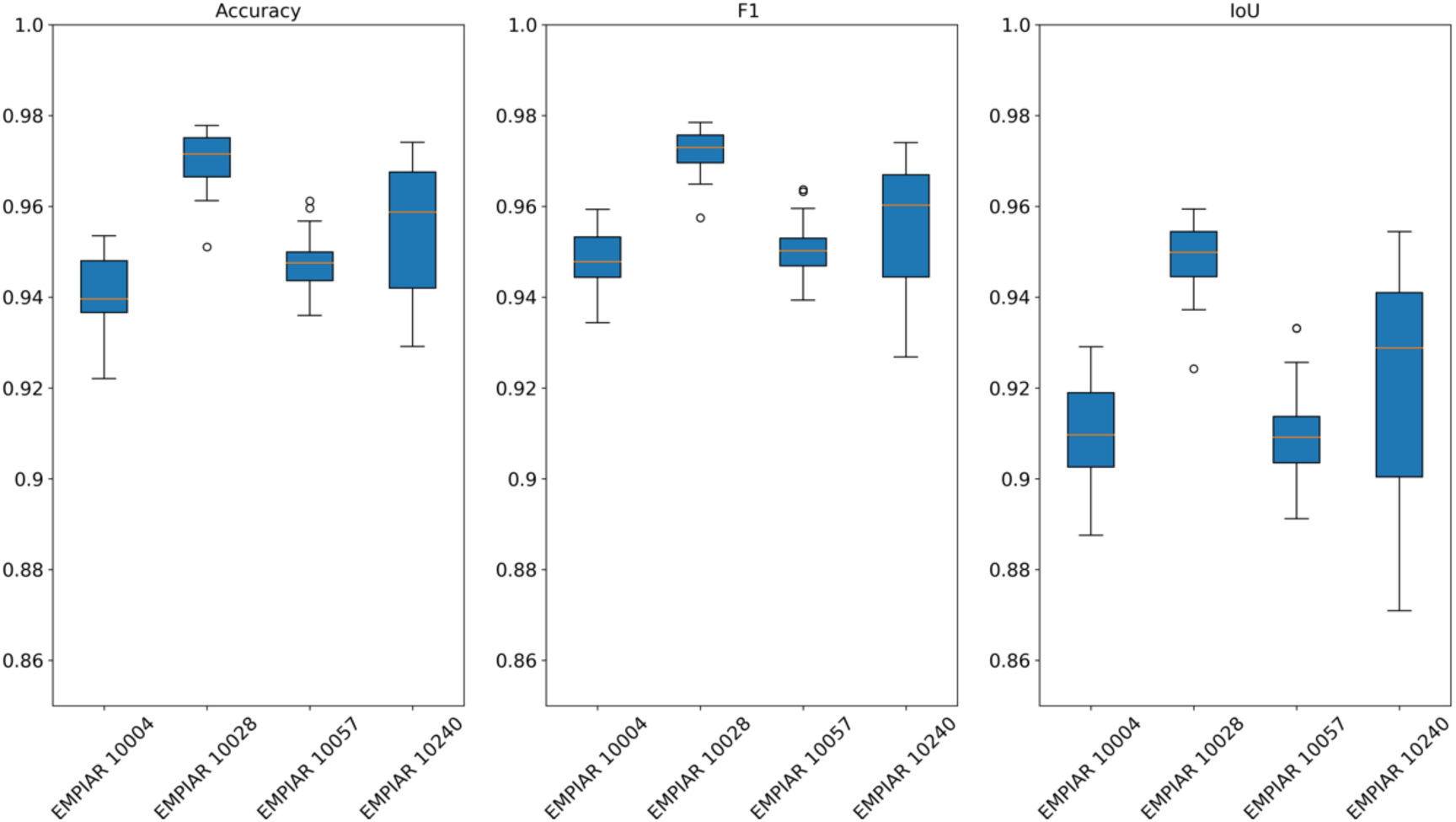
The boxplots showing the evaluation metrics of 12 micrographs from 4 proteins-**HIV-1 envelope glycoprotein** (EMPIAR 10004), **T20S proteasome** (EMPIAR 10057), **80S ribosome** (EMPIAR 10028) and **afTMEM16/nanodisc complex** (EMPIAR 10240). The box indicates the upper and lower quartiles and the median is shown as the line. The outliers and range of the data are represented by points and whiskers in the plot. The boxplots show the weighted average values of Accuracy, F1 and scores which are obtained by comparing the labels predicted using cross model with the ground truth labels.

## Discussion

In this study, a novel tool named CASSPER that can be used for automated particle picking from cryoEM images is presented. Using a powerful Semantic Segmentation Deep Learning framework, CASSPER colour codes all pixels in the cryo-EM micrographs to their probable classes. CASSPER is the first particle picking tool implementing the Residual Network architecture for efficient pixel-wise classification. Rather than searching for morphological features, it exploits the difference in the transmittance level of protein and non protein entities in the micrographs to locate the particles. This advantage translates into the ability of CASSPER to distinguish and pick the proteins in micrographs with ice contamination or carbon edges as shown in Figure 1.

CASSPER offers complete automation as it eliminates the need of manual particle picking at any stage in its operation. In the training phase, labels are generated using a GUI having four sliding bars to optimally adjust various filters as explained in the Methods section. It takes into account the intrinsic differences in the transmissivity of the particles in the image to generate colour coded labels for the particles in the micrographs. The particle coordinates are then extracted and returned in the popular star format for easy integration with any data processing software package such as RELION.

The advantage of learning the pixels that differentiate a protein from the rest of the image is that it can then be efficiently used for the discovery of the structures of new, unknown proteins. This is because in a Transmission Electron Microscope (TEM) image, what constitutes the image basically is the transmissivity of the media. If a machine can learn how each pixel, corresponding to the protein particle, differs from the rest, the collection of connected pixels can locate the position and shape of the protein. The method is only limited by the intrinsic differences in protein transmissivity that may cause fragmentation in the label for a single protein structure. Since this can be corrected visually, we allow the user to specify a size threshold based on the labels predicted by CASSPER before it is used to count and pick individual particles.

In earlier segmentation based tools like PIXER^27^, the feature map is segmented to get the protein containing regions and these regions are then given to a trained classifier to determine the particle centers. In CASSPER, the output of the SS network itself is classified into different classes in the micrograph and no additional post processing steps are needed for classification.

The prediction for TcdA1 and HCN1 using the CASSPER cross model trained on six other proteins yielded a resolution similar to the resolution achieved through the uniform pipeline. The promising results of evaluation metrics for unseen proteins using the cross model proves the capability of Semantic Segmentation network to distinguish the pixels based on their transmittance level.

## Methods

### Training Data Preparation

Semantic Segmentation is a very powerful tool to learn the relation between an object and its surroundings. The semantically segmented labels required for training the network is made using the CASSPER labelling tool. The processing steps of the tool is explained in **Figure 1**. The raw micrograph image is enhanced using a Gaussian filter followed by contrast enhancement. The combination of range and domain filtering in bilateral filter preserves the image edges and removes the noise in it. In our implementation, the filter size is tuned using the slide bars (**Figure 10** **(B)**). Sample vitrification usually leads to variable ice thickness in EM grid holes, this results in contrast difference in the micrographs. To overcome this situation, we employed a Contrast Limited Adaptive Histogram Equalization (CLAHE)^29^ on the images. The contrast limiting (CL) value for CLAHE can be adjusted using the slide bar. Usually low positive CL values are suitable for most proteins. The final segmentation in the enhanced image is done in two steps: a) Intensity thresholding and b) size filtering. Trackbars are used to set the intensity threshold value and the threshold value for the area of the particles. Based on these inputs, the final labels are created. This is a one time procedure and the generated values can be used for all the micrographs of the dataset under consideration, thereby, making the labeling procedure fast and accurate. In addition to protein particles, by following the same procedure, ice and carbon contaminants that have different transmittance are also labeled with different colors.

**Figure 10.**
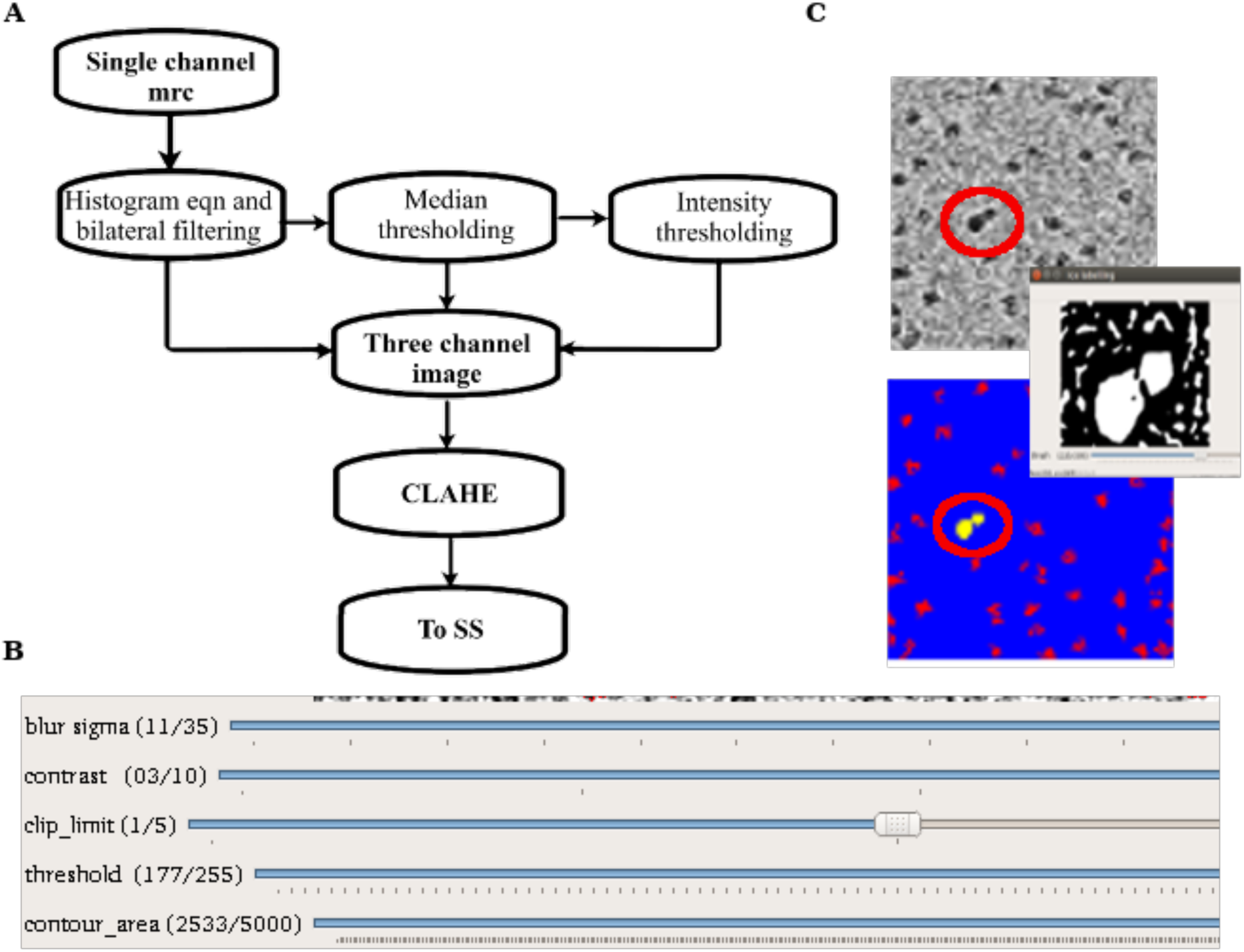
(A) The flowchart showing the pre-processing steps of CASSPER training phase. The single channel micrograph is enhanced and three different filters are applied to get the inputs for three channel image. **(B)** The toolbar section of the GUI which was used for labelling the proteins. **(C)** Figure showing the ice labelling method.

### Training of CASSPER

#### Preprocessing

The gray scale cryo-EM images have very low SNR hence, it has to be enhanced before submitting to the Semantic Segmentation algorithm. Also, the SS algorithm is designed to work on multichannel colour images. In our implementation, the three channel input image is obtained by applying three different filters to the motion corrected micrographs. Contrast enhancement and edge preserved noise removal of the input image are done by histogram equalization followed by bilateral filtering. This forms one channel. The second channel is prepared by Median thresholding that retains only pixels with intensities around a set threshold of the median pixel. In effect this channel enhances the contrast around the median pixel range of the image. The third channel of the image is generated by applying a Gaussian adaptive threshold to the second channel. These enhanced images are combined to form the three channel image. This image is finally adaptively histogram equalized using CLAHE to reduce the contrast difference effects and improve the efficiency of SS.

### Definition of the Statistical terms used

**True positive**-Number of pixels which are predicted to the correct class.

**False positive-** Number of pixels which are predicted to a wrong class.

**Precision-** Percentage of correct predictions.

**Recall-** Ratio of correct pixels in the predicted label to the ground truth.

**Intersection over Union (IoU)-** Ratio of the number of common pixels in the predicted and ground truth images to the union of the pixels in both images.

**F1 score**-Weighted average of precision and recall,

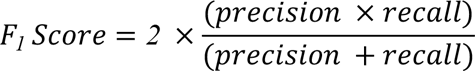

is used as the parameter for evaluating the validation performance and the model with highest F1 score is used for prediction of the unseen proteins.

For evaluating the performance of the prediction using the cross model, we employed F1, accuracy and mean IoU (Intersection over Union) scores to pixel-wise compare the predicted labels with the labels made using our labelling tool. The weighted average of particle and non particle pixels are indicated in these scores.

## Data Availability

The training datasets for this study, particle stacks and 2D class stacks are available on the GitHub page “CASSPER” along with a detailed practical manual for download under github page https://github.com/airis4d/CASSPER.

## Code availability

The source code is contained in the CASSPER software package and its use is restricted by the end user license agreement:Creative Commons License

## Acknowledgments

This work in J. Kumar lab is supported by the Wellcome Trust DBT India Alliance fellowship (grant number IA/I/13/2/501023). B. George thanks University Grants Commission and CMS College Kottayam for the Teacher Fellowship under FDP Scheme. A. Assaiya thanks DBT, India for a Senior Research Fellowship. We are also thankful to Dr. K. Vaghmare, and computational facility at IUCAA, Pune for helping with transfer and storage of raw EM data.

## Author Contributions

JK, RC, AKK, NSP and GP formulated the project and initiated the discussions at NCCS and IUCAA. The preprocessing and deep learning tools were developed and implemented by NSP, BG and RJR at airis4D. BG and RJR did testing and troubleshooting with inputs from AK, RC, AA and JK. The uniform pipeline, high-resolution 3D reconstruction and comparison with other tools were done by AA at NCCS with inputs from RC, NSP, AK and JK. The manuscript was read and approved by all the authors before submission.

